# Mapping and Analysis of QTL for Early Maturity Trait in Tetraploid Potato (*Solanum tuberosum* L.)

**DOI:** 10.1101/380501

**Authors:** Xingcui Li, Jianfei Xu, Shaoguang Duan, Jiaojiao Zhang, Chunsong Bian, Jun Hu, Guangcun Li, Liping Jin

## Abstract

Maturity is one of the important traits of potato. In order to get the genetic segment of potato early maturity trait, a tetraploid potato maturity segregation population of Zhongshu 19 × Zhongshu 3 was used for genetic analysis through the combination of high throughput simplified genome sequencing (2b-RAD) and bulked segregation analysis (BSA). A genetic segment related to the early maturity trait at the 3.7~4.2 Mb locus on the short arm of chromosome 5 was obtained and eight markers were developed based on this segment, while five of them were closely linked to the early maturity trait loci. Moreover, 42 SSR markers were developed based on the reference sequence of DM. Finally, a genetic map of chromosome 5 contained 50 markers was constructed using the Tetraploidmap software. The total map length was 172 cM with an average genetic distance of 3.44 cM. Combining with phenotypic data of the segregation population, we mapped the early maturity trait QTL with the contribution of 33.55% on the short arm of chromosome 5, located at 84cM between the flanking markers SSR5-85-1 and SCAR5-8 with the physical interval of 471kb. Gene annotation showed that there exist 34 genes in this region, 12 of them are unknown function. Among the other 22 annotated genes, E3 ubiquitin ligase gene *PUB14* may be related to maturity and regulate tuber formation. Our fine mapping of the early maturity QTL made a solid foundation for cloning of the early maturity controlled gene or genes.

**Key message:** Early maturity site was mapped using a tetraploid potato segregation population derived from cv. Zhongshu 19 and Zhongshu 3. One major QTL with 33.55% contribution to early maturity was fine mapped in physical interval of 471kb on chromosome 5.

## Introduction

Potato (*Solanum tuberosum* L.) is the third largest food crop worldwide and also serves as an important industrial raw material. Maturity is one of the important features for identifying the characteristics of potato varieties. It is also an important agronomic trait and a major breeding target. Different maturity varieties can meet the production and consumption demand in different regions and seasons. The breeding of different maturity varieties is greatly significant in the development of the potato industry. Previous studies showed that potato maturity traits were controlled by minor polygenes and were recessive, and the potato maturity quantitative trait loci (QTLs) were distributed on 12 chromosomes of potato. However, most of the studies have mapped the major QTL for maturity on chromosome 5 and linked to the QTL for resistance to late blight (Van Eck et al. 1995; Collins et al. 1999; Oberhagemann et al. 1999; Ewing et al. 2000; Draffehn et al. 2013). In 2007, Sliwka et al. (2007) used a diploid potato population to map a QTL associated with growth stages and closely linked to molecular marker BA47f2t7 (P1) on chromosome 5 (Sliwka et al. 2007). Later, scientist found that the major QTLs that control maturity on chromosome 5 were closely related to the QTLs controlling late blight resistance but were independent of each other and closely linked to molecular marker GP21 (Danan et al. 2011). Using tetraploid materials, a major QTL was also obtained to be associated with maturity on chromosome 5, which showed a contribution rate of 54.7% (Bradshaw et al. 2004). Later, by increasing the number of progeny populations, some minor QTLs with contribution rates between 5.4% and 16.5% were identified (Bradshaw et al. 2008). In recent years, a large number of single nucleotide polymorphisms (SNPs) have been developed, and a high-density genetic linkage map containing 3,839 SNPs has been constructed by analyzing dose information of SNPs in tetraploid potato. One QTL associated with maturity closely linked to molecular marker c2_476095 has also been mapped on chromosome 5, with a contributing rate of 55% (Hackett et al. 2014). In 2013, Kloosterman et al. (2013) through fine mapping cloned the *StCDF1.2* gene related to potato maturity near molecular marker GP21 on the short-arm of chromosome 5. This gene belongs to the DNA-binding One Zinc Finger transcription factor family, which involves in tuber formation as an intermediate regulatory factor in the potato tuber induction pathway.

Tetraploid potato (2n = 4x = 48) is highly heterozygous, with high frequency of genetic recombination and severe recession after self-fertilization (Manrique-Carpintero et al. 2018). The molecular marker is difficult to develop. The diploid potato chromosome number is small, the genetic background is relatively simple, and it is easy to carry out genetic analysis and operation. Therefore, the development of potato markers and genetic map construction is mainly at the diploid level, and is done mainly with the use of the traditional QTL mapping analysis method, whereas fewer studies are conducted on tetraploid materials. Although most of the studies have mapped QTLs on chromosome 5, there are no more genes related to potato physiology maturity are cloned except for *StCDF1.2* gene.

With the development of high-throughput sequencing technology, marker development and genetic segment mining in complex genomic species becomes simple. The high-throughput simplified genome sequencing 2b-RAD technique uses DNA type II B restriction endonucleases *(BsaXI* and *AlfI)* to cleave DNA from the upstream and downstream sites of the target site on genomic DNA to obtain DNA fragments of consistent length (tags). The 2b-RAD technique avoids the process of fragment size selection in other genome-wide sequencing technologies, making tags more uniform in the genome, allowing for the development of the complex genomic marker, mining of chromosomal segments, and construction of linkage map and genetic variation map in natural populations (Wang et al. 2012). At present, this technique has been applied to construct high-density genetic linkage maps of animals, plants, and marine organisms. The high-density genetic linkage map of Chlamys Farrer includes 3,806 molecular markers, the average distance between markers is 0.41 cM with genome coverage of 99.5%, and is it mapped to growth-and-sex-related QTLs (Jiao et al. 2013, 2014).

In this study, we intend to use the tetraploid potato maturity segregation population, combine high-throughput simplified genome sequencing 2b-RAD with the BSA method, to identify the genetic segment related to the early maturity trait and develop molecular markers based on the genetic segment. Linkage Map was constructed by using software TetraploidMap for Windows (Hackett et al. 2007), which was designed for calculating linkage maps from the maker phenotypes of the parents and segregating progeny of a cross in an autotetraploid species. Combined with 4 years foliage maturity phenotypic data, QTL analysis was performed using this software, which will provide a foundation for cloning and functional verification of the maturity gene or genes.

## Materials and methods

### Plant materials

The mapping population, consisted of 221 individuals, was from a cross between two tetraploid varieties: ‘Zhongshu19’ × ‘Zhongshu 3’. The female parent ‘Zhongshu 19’ with a pedigree of (CIP92.187×CIP93.154) × (Bierma × Colmo) is a late maturing variety with a growth period of 110 days after emergence, and the male parent ‘Zhongshu 3’ with a pedigree of (Jingfeng 1 ×BF77A) is an early maturing variety with a growth period of 75 days. They were bred by the Institute of Vegetables and Flowers, Chinese Academy of Agricultural Sciences, Beijing.

### Field tests

During four consecutive years (2014-2017), the field experiment was carried out in Zhangbei (ZB) County of Hebei Province (41°15’ N, 114°07’ E, 1500 m a.s.l.) and Liaocheng(LC) City of Shandong Province (36°45’ N, 115°97’ E, 38 m a.s.l.) in China. The mean daily temperature ranged from 10.05 °C to 21.66 °C and 12.59 °C to 24.37 °C corresponding to the average minimum and maximum temperatures. The average sunshine duration is 8.54 h and 6.45 h respectively. Four tubers of each progeny were sown in early May and mid-March and harvested in late September and late June in ZB and LC respectively in each year.

### Phenotypic identification

Growth period was defined the time from emergence (seedling emergence) to physiological maturation (50% of plant leaves appear yellow) in this study. Individual emergence was investigated every five days from 20 days after sowing. Physiological maturation time of the individuals was investigated every five days from 60 days after emergence (DAE), continued to investigate every five days, and finally calculated the growth period of each individual.

The maturity of the F1 population was divided into five groups according to the growth period, and scored from one to five scales. 1: very early mature type with a growth period less than 70 DAE; 2: early mature type with a growth period 71-80 DAE; 3: middle mature type with a growth period 81-100 DAE; 4: late maturity type with a growth period 101-110 DAE; 5: very late mature type with a growth period more than 110 DAE.

### DNA extraction and progeny mixed pool construction

Two-hundred twenty-one samples were collected from young leaves of Zhongshu 3, Zhongshu 19, and F1 plants. The genomic DNA was extracted by the CTAB method, and DNA quality was detected with 1% agarose gel and BioDrop for marker development and validation.

We used a punch to take one young leaf from the same part of each F1 generation early and late maturing material with a growth period of less than 70 DAE and greater than 110 DAE, respectively, mixed them in equal amount, extracted their DNA, and constructed an early and late maturing mixed DNA pool for simplified genome sequencing. The DNA extraction method and quality testing method are the same as mentioned above.

### Simplified genome sequencing analysis and marker development

A high-throughput simplified genomic 2b-RAD sequencing technique was used to construct a tag sequencing library of parental DNA and extreme progeny DNA pool. The single-terminal sequencing was performed on the Hiseq 2500 v2 platform. After removing from original reads of the sequences that contain no *BsaXI* recognition sites, low quality sequences and sequences with more than 10 consecutive identical bases, the individual high quality reads were mapped to the potato DM reference sequence using SOAP software. The number and depth of specific tags that can be used for typing were obtained, and genome-wide SNP screening and typing was conducted. According to the classification results, we constructed a chromosome tag density distribution map of four samples with Matlab software, further, constructed specific tag density distribution and absolute value distribution of the specific tag density between the two mixed pools, respectively, and then screened out chromosome segments with a large difference in tag densities.

Based on the tag information obtained in the different segments, the synthetic primers were designed according to the genomic sequences on the potato genome sequence website (http://solanaceae.plantbiology.msu.edu/cgi-bin/gbrowse/potato/). The quality and specificity of the primers were detected by using the genomic DNAs of the late maturing parent Zhongshu 19 and early maturing parent Zhongshu 3 as the templates. The amplified products were detected by agarose gel electrophoresis with a concentration of 1.2%. The PCR reaction system consisted of: 5.6 μL ddH_2_O, 1.0 μL buffer (10 × PCR), 0.8 μL dNTPs (10 mmol L^−1^), 0.2 μL forward primer (10 μmol L^−1^), 0.2 μL reverse primer (10 μmol L^−1^), 0.2 μL Taq enzyme (2.5 U μL^−1^), and 2 μL DNA (25 ng μL^−1^). PCR was performed as follows: 94°C for 3 min, followed by 35 cycles of 94°C for 30 s, 59°C for 30 s, and 72°C for 50 s, and finally, 72°C for 10 min.

The PCR products were digested by *BsaXI* restriction endonuclease if the primers could amplify clear and non-different bands in the parents (Zhongshu 3 and Zhongshu 19). The enzyme digestion system was 8 μL PCR product, 1.5 μL CutSmart® buffer, 0.2 μL *BsaXI* endonuclease (0.5 μL/U), and 5.3 μL ddH_2_O. The primers whose PCR products can be digested only in one parent and not digested in the other parent were used to develop CAPs markers. The primers that amplified a band in one parent but not in the other parent, or amplified different size bands in the two parents, were used to develop SCAR markers. Finally, the progeny early and late maturing materials were used for validation.

### Genetic map construction and QTL mapping

Polymorphism screening of the molecular markers developed in this study and the previously published 152 SSR markers on chromosome 5, was conducted using the genomic DNAs of Zhongshu 3, Zhongshu 19, and 16 F1 generation materials as the templates (Feingold et al. 2005; Ghislain et al. 2009; Zhou, 2014), and the selected polymorphic markers were verified using the population of 221 progenies. According to the requirement of tetraploid mapping software TetraploidMap (Hackett et al. 2007), we selected the following types of makers for map construction: (1) simplex dominant markers (segregating 1:1) with a *p* value greater than 0.001 from a Chi-square test for goodness of fit; (2) duplex dominant makers (segregating 5:1)with a *p* value greater than 0.01; (3) double-simplex dominant makers (segregating 3:1) with a *p* value generally greater than 0.01 and which were known to be linked to at least one simplex marker; While dominant makers present in both parents are extremely uninformative about recombination unless the segregation ratio is 3:1.(Hackett at al. 2007). So makers with the segregation ratio is 11:1 (simplex × duplex), were in fact omitted from the linkage map. Because of all makers we chose all from chromosome 5, so most of makers were selected to construct linkage map for chromosome 5, except for a few makers that may belong different linkage group since add them the software would show an error in subsequent QTL analysis. After the makers are selected, the TetraploidMap ordered them for proper sequence and then we assigned 50 makers to four homologous chromosomes considering with the relationship between makers is couple or repulsive (Simonsen and McIntyre. 2004). The primer information for makers in this research are listed in Table 1. And genotypes information for 221 progenies are listed in Table S1. Consequently, combination with the mean value of 4 years progeny phenotype data, details see Table S2. the maturity QTL was mapped by using TetraploidMap.

### Data availability

The authors state that all data necessary for confirming the conclusions presented in the manuscript are represented fully within the manuscript and supplemental information at Figshare: https://figshare.com/s/876e1539f3f42b8ec8ef. TetraploidMap for Windows is freely available at: http://www.bioss.ac.uk/ (user-friendly software).

## Results

### Distribution of the segregation population maturity

In ZB, the materials were planted on 20^th^ May and emerged in mid-June. Part of the genotypes reached physiological maturity in late August, while others were still keep vigorous in late September. In LC, the corresponding date above were 10^th^ March, mid-April and late June respectively. According to the field investigation in the four consecutive years, the statistical analysis of the population for maturity trait was consistent with polygenic genetic characteristic of quantitative traits. The offspring showed a normal distribution over the entire scoring range (Fig. 1). Detail phenotypic data among four years was shown in Table S1. Genomic DNA was extracted from 35 very early and early maturing genotypes and 33 very late maturing genotypes, and the corresponding early maturing DNA pool and late maturing DNA pool were constructed.

**Fig. 1.**
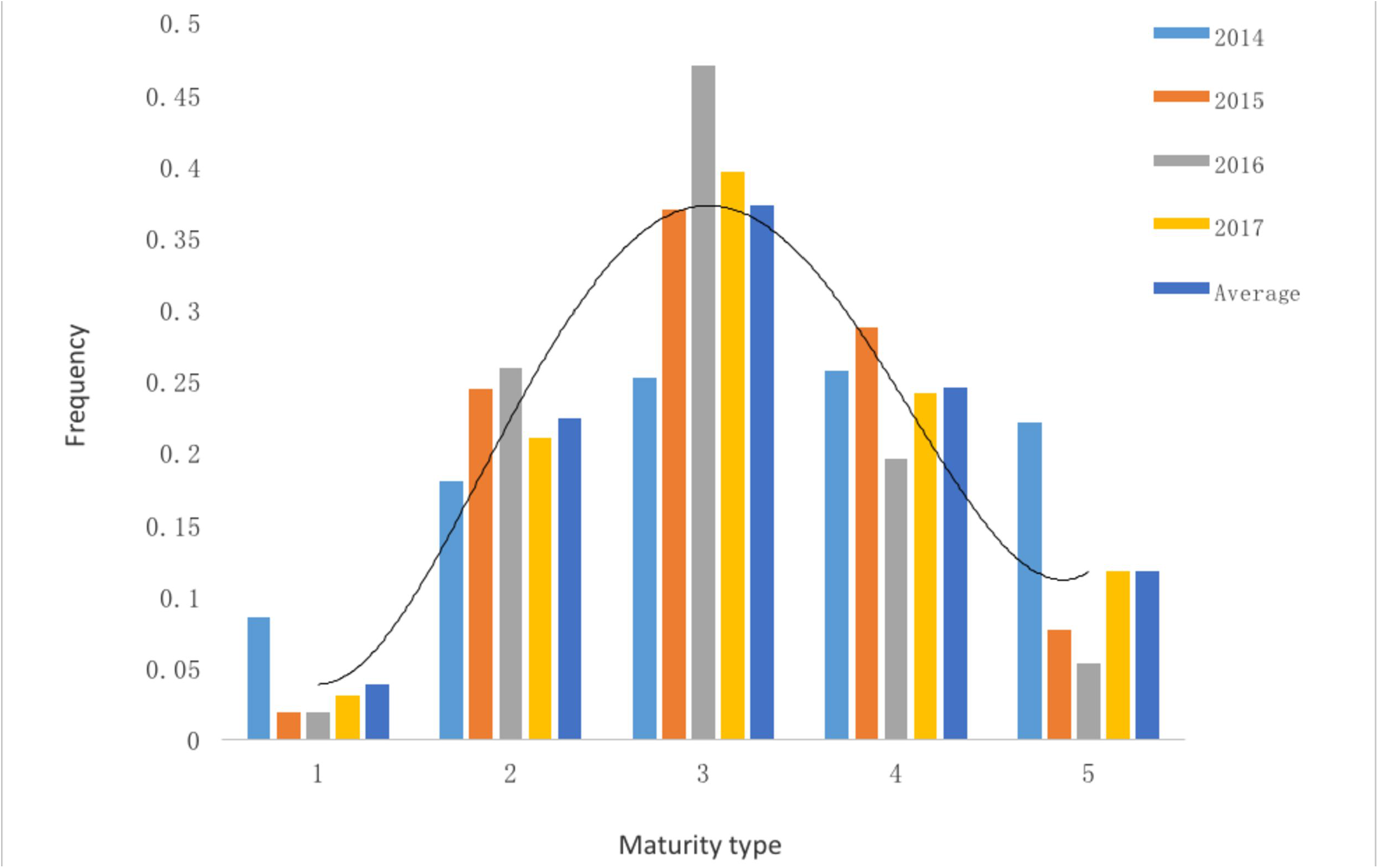
Maturity frequency distribution of the mapping population individuals. X-axis indicates maturity type of the mapping population, Y-axis indicates the ratio between individuals of different maturity types and the whole mapping population. The entire mapping population consists of 221 individuals.

### Screening and marker development of early maturity traits

Simplified genome sequencing on four samples of parents Zhongshu 3 and Zhongshu 19, the early maturing pool, and the late maturing pool were carried out using the 2b-RAD technique, and a total of 52,714,218 reads were obtained. The average number of reads of each sample was 13,178,554 with an average sequencing depth of 42× (Table 2). High-quality reads containing *BsaXI* digestion sites in the four sequencing libraries were all higher than 90%, and finally 125,556 unique tags in average of each sample were obtained.

SNP marker typing was performed according to the obtained sequencing tag sequences and the tag distribution and density profiles on 12 potato chromosomes were got, and the results showed that the tags were evenly distributed on all 12 potato chromosomes, and there was no large-scale tag information missing (Fig. 2). Furthermore, according to the specific tag density of early maturing and late maturing pools, the density difference map of the specific tag between early maturing and late maturing pools was obtained, and the results showed that the segments with a large difference in tag density between early and late maturing pools were mainly concentrated on chromosomes 4 and 5, where in the segment with the most significant difference in tag density was found at 3.68–6.19 Mb on chromosome 5, followed by 18.6–20.9 Mb and 27.4–30.3 Mb on chromosome 4 (Fig. 3). Moreover, the specific tag density of the early maturing pool among the differential segments of chromosome 5 was relatively high, suggesting that this segment may be related to the early maturity trait. But all the specific tag densities of the late maturing pool were relatively high among the differential segments of chromosome 4, which might be related to the late maturity trait (Fig. 3).

**Fig. 2.**
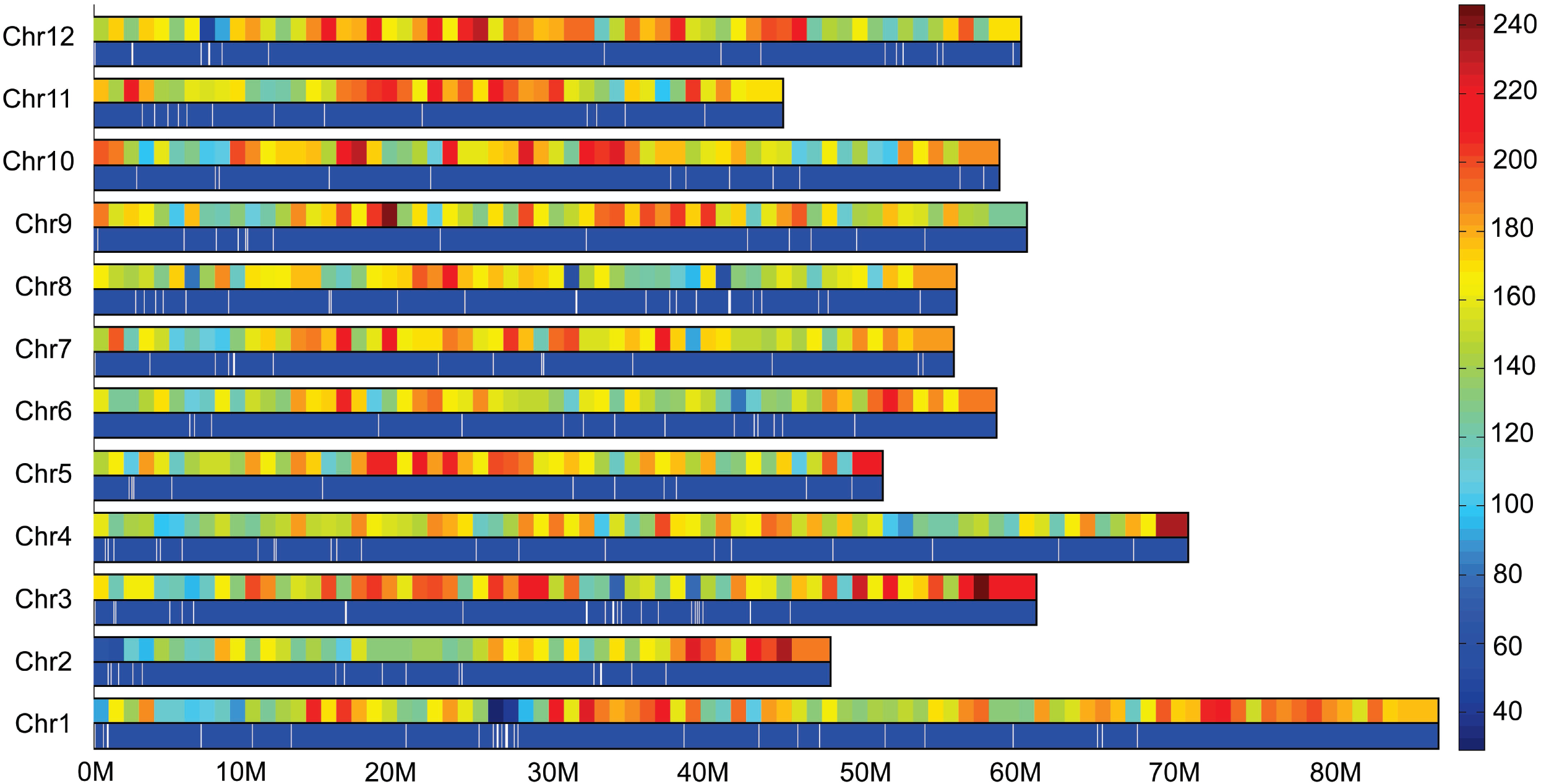
Distribution of tags on the chromosome and tag density map. 0-80 M: chromosome physical distance (unit: M). Chr1-Chr12: 12 potato chromosomes, respectively. Right histogram: different colors indicate different tag numbers, and numbers indicate specific tag numbers. In each chromosome, the upper long histogram shows the tag density distribution on the chromosome, and the long histogram below shows the tag distribution on the chromosome.

**Fig. 3.**
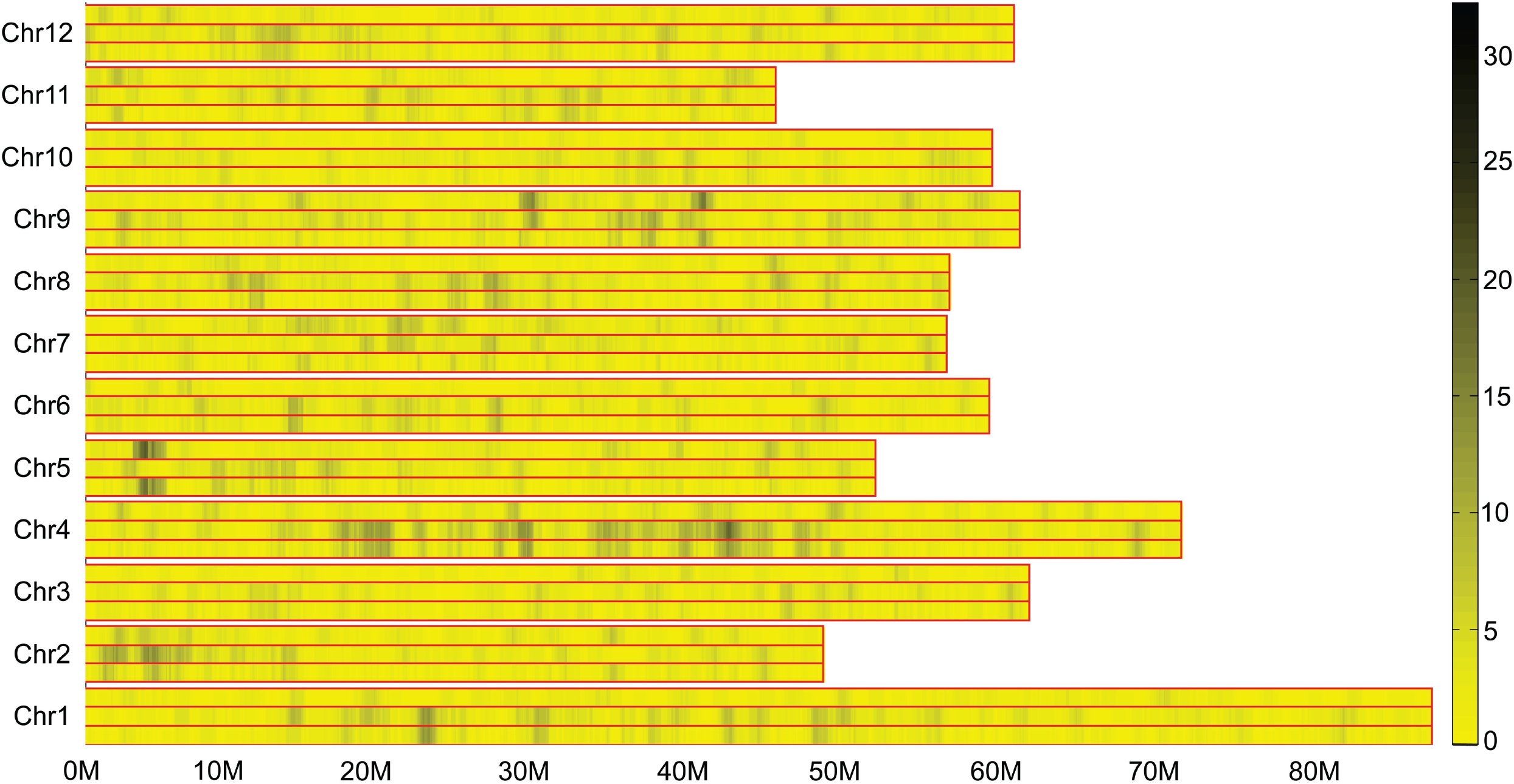
Differential tag density map of early and late maturing mixed pool. 0-80 M: chromosome physical distance (unit: M). Chr1-Chr12: 12 potato chromosomes, respectively. In each chromosome, the upper long histogram shows the specific tag density of the early maturing pool, the middle long histogram shows the specific tag density of the late maturing pool, and the long histogram below shows the absolute value of differential tag density between the two pools. Right histogram, different colors indicate different tag numbers, and numbers indicate the specific tag numbers.

Based on the above specific tag positions of chromosomal differential segments, combined with potato reference genomic sequences, a total of 92 primers were designed with 52 pairs of primers in the 3.68–6.19 Mb region of chromosome 5 and 40 pairs of primers in the differential segments of chromosome 4. Twenty polymorphic markers including 4 CAPS and 4 SCAR markers located on chromosome 5 were obtained from 92 designed primers. Further test on the F1 individual plants showed that only five markers of SCAR5-5, SCAR5-8, CAPS5-3-2, CAPS5-21-2 and CAPS5-24 were closely linked to the early maturity trait. Polymorphic markers on other chromosomes were not linked with maturity trait (data not shown), suggesting that only the genetic segment on chromosome 5 was associated with the early maturity trait. The marker SCAR5-8 on chromosome 5 was used to validate 70 early and late maturing Chinese cultivars, and the results showed that the coincidence rate of marker detection results and trait identification results was as high as 81.4% (Li et al., 2017), indicating that this marker could be applied in potato molecular marker-assisted breeding.

### QTL mapping for early maturity trait

In order to verify the accuracy of genetic segments for the early maturity trait that were mined, we constructed a genetic linkage map of chromosome 5 and performed QTL mapping for the early maturity trait. First, early and late maturing parents and their 16 progenies were used to test the polymorphism of the 152 pairs of SSR primers on chromosome 5, and a total of 32 polymorphic SSR markers were obtained. Then the 32 polymorphic SSR markers and the previous eight molecular markers (4 CAPS, 4 SCAR markers) developed on chromosome 5 were further used to test the F1 generation. In this study, we focused on early maturity, and the genetic linkage map for early type parent ‘zhongshu 3’ was constructed (Fig 4). The map including a total of 50 makers, of which 33 were simplex, 4 were duplex, and 13 were double simplex makers (TetraploidMap generate automatically combined with the segregation ratio) (Table 3). The total length of the map coverage was 172 cM, and the average distance between markers was 3.44 cM. The 50 markers of this map were four CAPS markers, four SCAR markers, and 42 SSR markers, respectively. 42 SSR markers were consisted of 25 single markers and 17 polymorphic SSR markers. The 25 single markers were come from the division of 11 polymorphic SSR markers, and they were SSR5-22-1, SSR5-22-2, SSR5-36-1, SSR5-36-2, SSR5-38-1, SSR5-38-2, SSR5-40-1, SSR5-40-2, SSR5-40-3, SSR5-55-1, SSR5-55-2, SSR5-85-1, SSR5-85-2, SSR5-85-3, SSR5-85-4, SSR5-100-1, SSR5-100-2, SSR5-103-3, SSR5-103-4, PM0333-2, PM0333-3, STI049-2, STI049-3, STG0021-1, and STG0021-2. Actually, among the 32 polymorphic SSR markers only 28 markers were used to construct the genetic linkage map.

**Fig. 4.**
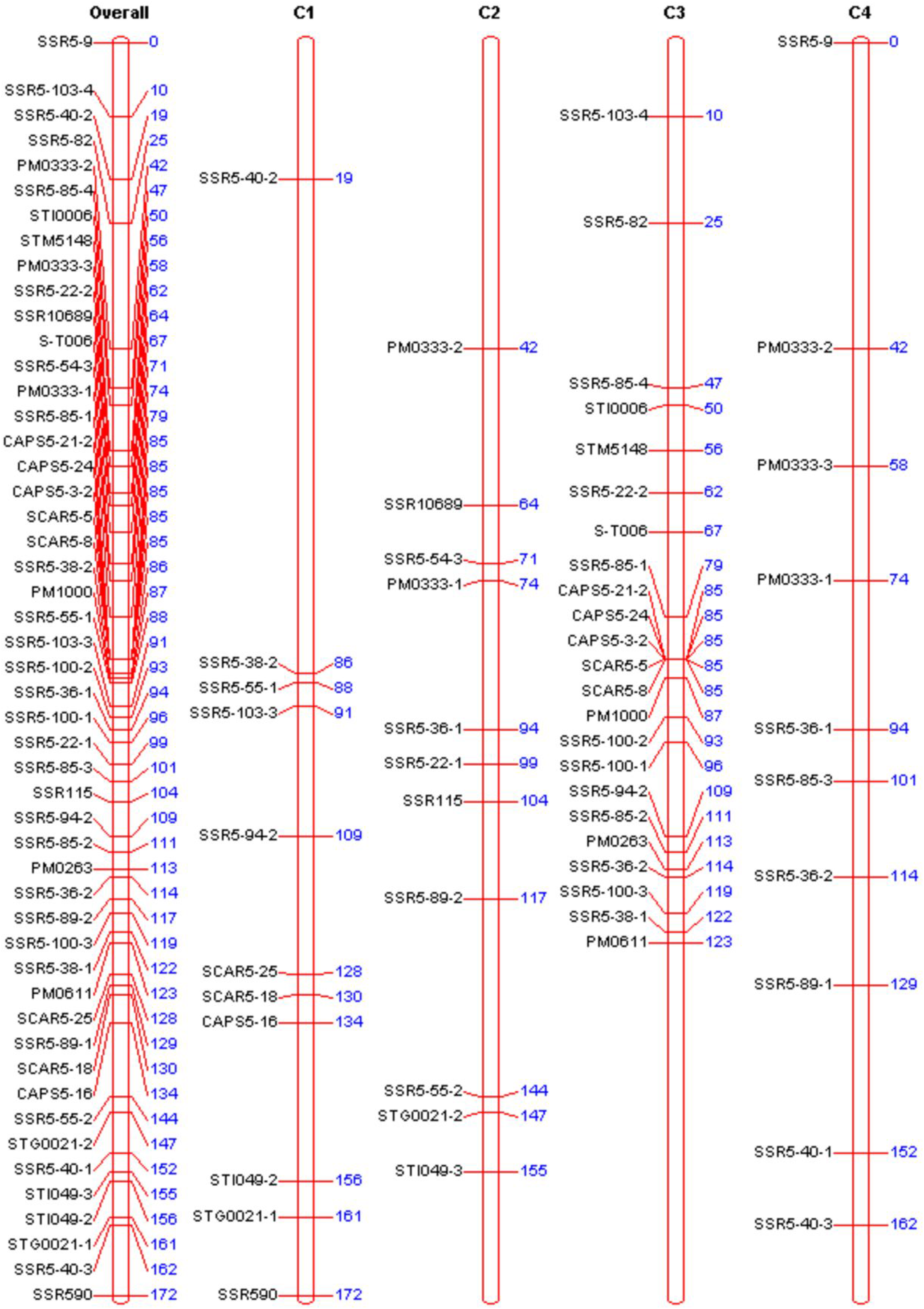
Genetic map of chromosome 5 in potato. The left side of the map is 50 molecular markers. The numbers on the right indicate the corresponding genetic distances (unit: cM). Overall: chromosome 5. C1-C4: represent four homologous chromosomes of chromosome 5, respectively.

QTLs for foliage maturity type were identified using TetraploidMap. Initially, the significance of individual markers for each year trait was tested by analysis of variance (ANOVA) and the analysis procedure Kruskall-Wallis in TetraploidMap. This two tests were used to compare the differences of the mean value of different makers genotype (Simonsen and McIntyre, 2004). A P-value of less than 0.01 was used as a threshold criterion for QTL detection. Results from above two tests suggested the existence of QTLs for foliage mature type on chromosome 5. There are 22 makers on chromosome 5 may associated with maturity. The maker cluster including maker CAPS 5-21-2, CAPS5-24, CAPS5-3-2, SCAR5-5, SCAR5-8 were closely linked to the QTL locus, since not only their P value are 0.000 but also their smallest SED with 0.0937 (Table 3).

Furthermore, in combination with phenotypic identification data of F1 segregation population, three significant QTLs for early-maturity trait loci were identified with LOD scores above the threshold value of 2.97 (The blue dotted line) located on chromosome 5 (Figure 5). The most remarkable QTL was mapped at 84 cM of the 3rd homologous chromosome of chromosome 5 between marker SSR5-85-1 and maker cluster including CAPS5-21-2, CAPS5-24, CAPS5-3-2, SCAR5-5, SCAR5-8, but more closely linked to the 5 markers, which was in line with the position of the maturity genetic segment mined in this study. The maximum LOD value of this QTL was 19.491 explaining 35.55% of the phenotypic variation. And there were other two locus may also associated with maturity type, one QTL linked to marker STM5148 with the LOD value 12.50, another one mapped at 113cM in correspondence of marker PM0263 with the LOD value nearly 11.30.

**Fig. 5.**
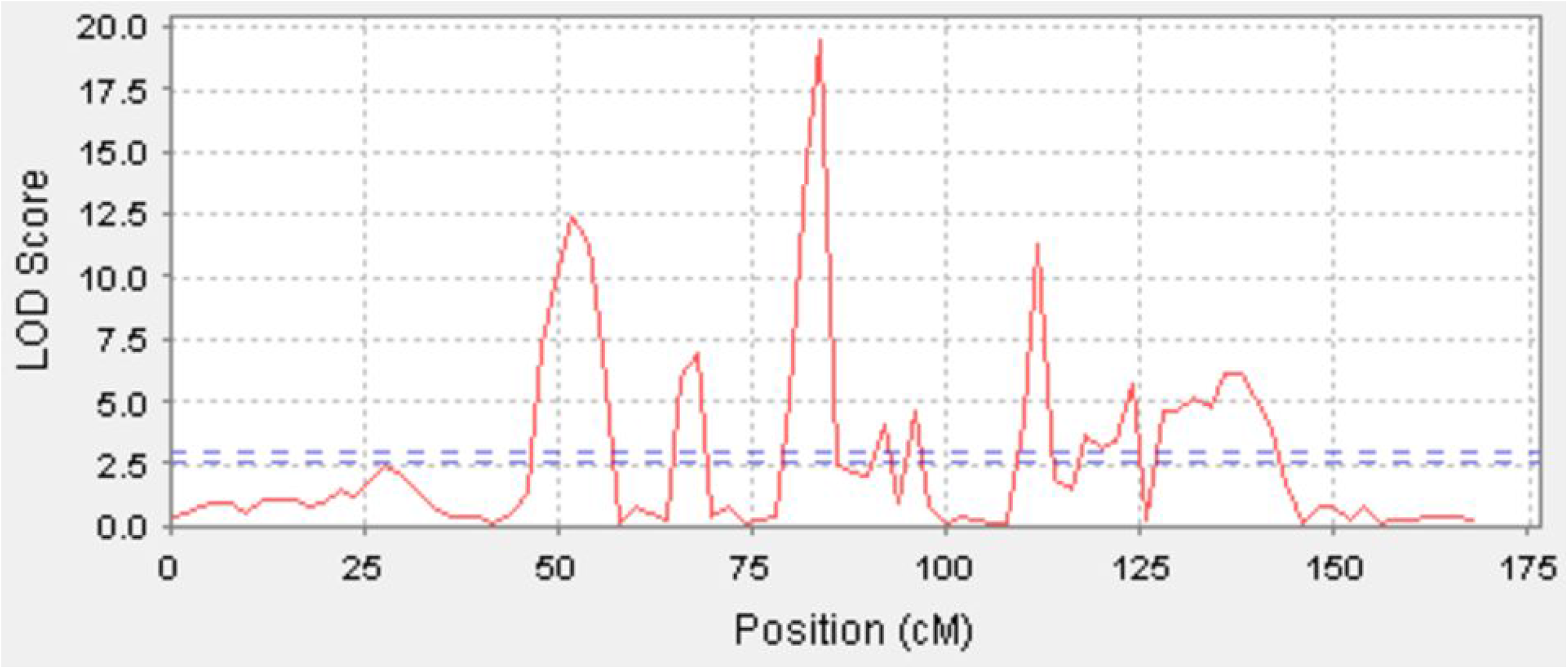
QTL mapping for Maturity. Line shows LOD values over the chromosome 5. The horizontal line indicates the confidence threshold (LOD value =2.97).

In conjunction with the marker locus and reference genome of the DM sequence, the QTL of early maturity trait was mapped to the 471 Kb physical interval between the flanking markers of SSR5-85-1 and SCAR5-8, which contained six molecular markers (Fig. 6). The order of all makers in this QTL region is consistent with the physical order of the makers in the potato DM genome sequence. (PGSC. Tuberosum group Phureja DM1-3 Pseudomolecules (v4.03)). Gene prediction and annotation showed that there are 34 genes in this 471 Kb physical range based on the comparison with the reference genome of DM sequence, (http://solanaceae.plantbiology.msu.edu/pgsc_download.shtml), of which 22 genes have been annotated and 12 genes functions are unknown (Table 4). The 22 annotated genes have different physiological functions and participate in different physiological regulation pathways. For example, E3 ubiquitin ligase PUB14 is involved in the photoperiodic regulatory pathway. The auxin export carrier is involved in plant hormone transport. Heat shock protein binding, Quinolinate phosphoribosyl transferase, phosphatidylinositol kinase fyv1, resistance protein BS2, and resistance protein PSH-RGH6 are involved in plant defensive reactions. The WD repeat protein is involved in protein transport and nucleic acid processing modification. The Myb transcription factor is involved in the plant secondary metabolic regulatory pathway.

**Fig. 6.**
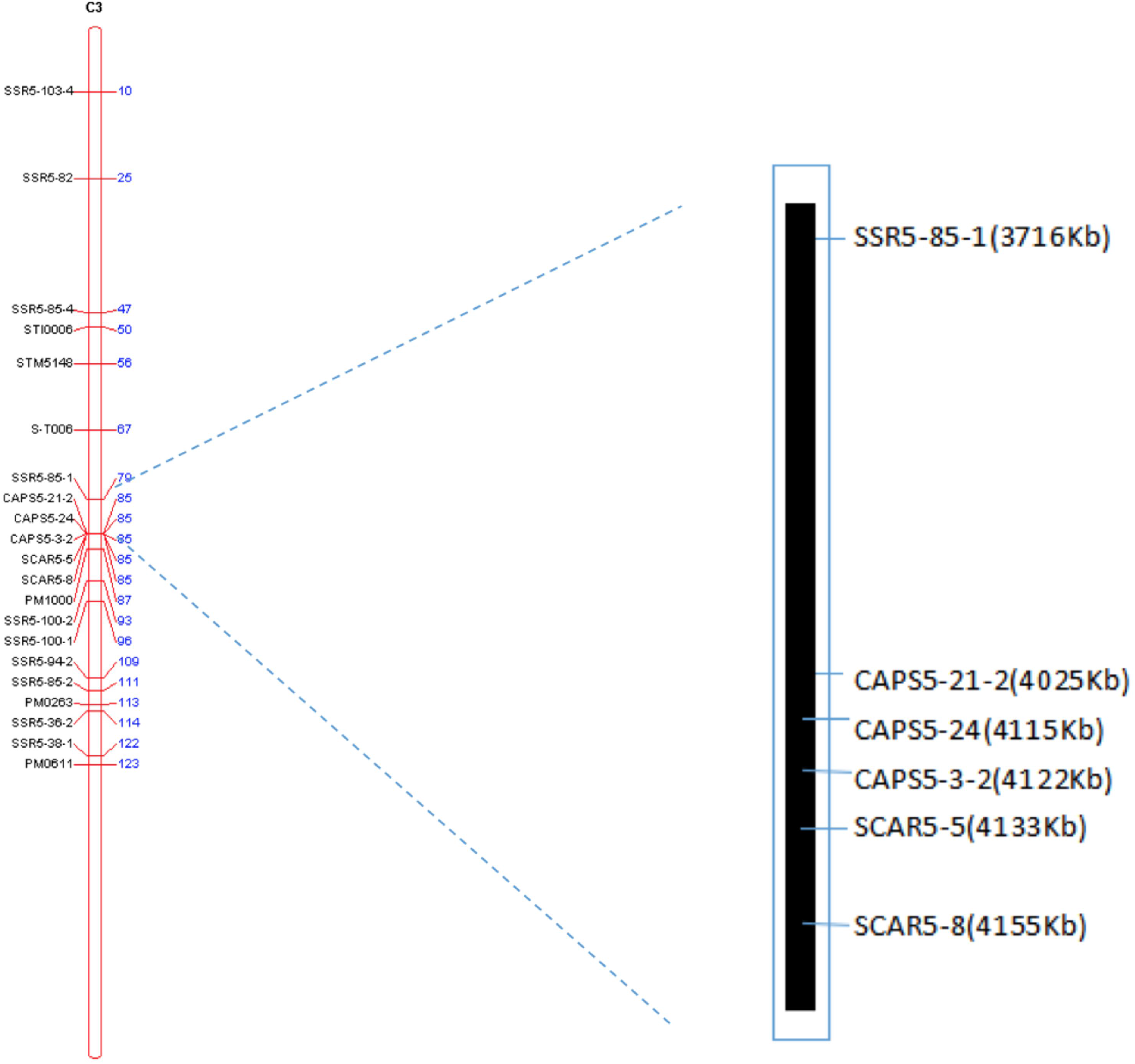
Physical interval of the early maturity QTL on chromosome 5. The left side is genetic linkage map of 3^rd^ homologous chromosome 5, and the right one is corresponding physical map.

**Table 4.**
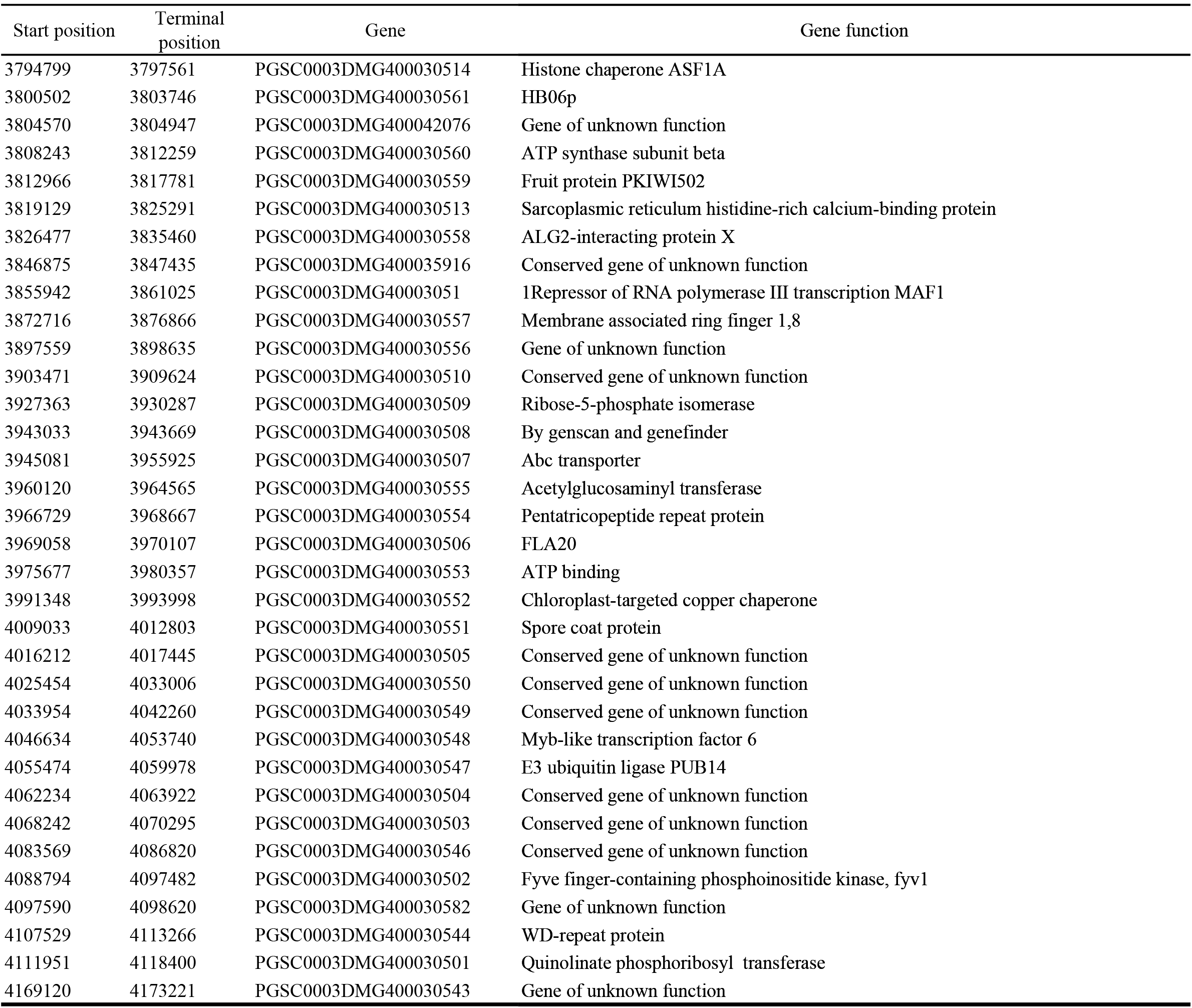
Gene annotaition and location in the physical interval

## Discussion

### Genetic segments mining for early maturity trait based on 2b-RAD simplified genome sequencing

At present, many researchers have used traditional molecular marker technologies to carry out potato QTL mapping. Most of the studies mapped maturity major genetic loci on chromosome 5 and found that other chromosomes may also have maturity micro-genetic sites. In this study, for the first time, we used the high-throughput simplified 2b-RAD sequencing technique to identify and mark genetic segments for potato complex quantitative traits, found a segment with large difference in specific tags on chromosome 4 and 5, and further tag validation showed that the genetic segment on chromosome 5 is associated with the early maturity trait, which is consistent with previous studies (Collins et al. 1999; Oberhagemann et al. 1999; Ewing et al. 2000; Bradshaw et al. 2004, 2008; Danan et al. 2011; Hackett et al. 2014).

Tetraploid potato has a complex genetics, and molecular marker development is difficult. Previously developed markers were mainly based on diploid (Bakker et al. 2004; Asano et al. 2012; Zhu et al. 2015; Hara-Skrzypiec et al. 2018), but markers developed directly at the tetraploid level were rarely reported. This study based on the maturity-related genetic segments mined on chromosome 5, a total of 52 pairs of specific primers were designed and synthesized, and eight molecular markers were developed with a development efficiency of 15.4%. The eight markers were further verified by early and late maturing progenies in the tetraploid segregation population, five markers were found to be closely linked to the early maturity trait. Therefore, the method can be used for the development of markers of complex genomes such as potato. Combined with published potato reference genomes, this method can greatly shorten the marker development cycle and map the markers. Which will also provide a reference for marker development of other polyploid complex genomic organisms.

In this study, we found that not only was there a maturity related genetic segment on chromosome 5, but there were also some segments with large difference in specific tag density on chromosome 4. Although there are a few reports that mention the existence of a maturity-related genetic segment on the chromosome 4, the tag validation results in this study showed that the genetic segment on chromosome 4 was not related to maturity. This may be due to several reasons: First of all, the tetraploid potato genome is highly heterozygous, more complicated than diploid potato in heredity, and in the high-throughput sequencing process, may be due to sequencing depth not being enough or *BsaXI* recognition sequence preferences; some information was not measured. Secondly, due to the filtering out some of the low quality reads resulted in partial loss of genetic information, and eventually the differential tag interval appears. Thirdly, when performing specific tag filtering, removing the tags with fewer occurrences may also cause a difference in tag density.

### Genetic linkage map construction and QTL mapping

In this study, a 50-marker genetic linkage map was constructed using tetraploid potato segregation population, and an early maturity trait QTL was mapped at the position of 84 cM near the marker SCAR5-8 (or SCAR5-5, CAPS5-3-2, CAPS5-24, CAPS5-21-2) on the short arm of chromosome 5. Which indicates that the QTL for the early-maturity trait obtained by genetic map mapping is consistent with the maturity genetic segment results obtained by high-throughput simplified genome sequencing, and also proved the feasibility and accuracy of high-throughput simplified genome sequencing 2b-RAD method in mining potato genetic segments for important traits. The five molecular markers (SCAR5-8, SCAR5-5, CAPS5-3-2, CAPS5-24, CAPS5-21-25) that are closely linked to the early maturity trait loci are from the same genetic segment and are relatively close to each other, and no significant separations between the markers in the genetic map. Furthermore, the five molecular markers were clustered on the 3rd homologous chromosome and belonged to the same linkage group. The other three molecular markers, CAPS5-16, SCAR5-18, SCAR5-25 probably linked to the late maturity trait loci, were clustered on the 1st homologous chromosome, and belonged to another linkage group (Fig. 4).

Previous studies on the maturity QTL mapping showed that the genetic interval of maturity QTL was relatively large. The major QTL for maturity was mapped at 0–6 cM on chromosome 5 in diploid materials, and the genetic distance between two flanking markers was 6 cM (Visker et al. 2003). In tetraploid materials, the major QTL for maturity was also mapped at 14–22 cM on chromosome 5, and the genetic distance between two flanking markers was 8 cM (Hackett et al. 2014). In this study, based on simplified genome genetic segment mining and marker development, the QTL for the early maturity trait was mapped in a physical interval of 471 kb, which greatly approached the early maturity trait loci and the average distance between two makers was 3.44 cM. Bioinformatics analysis revealed that there were a total of 22 annotated genes in the 471kb region, and the E3 ubiquitin ligase PUB14 gene was most likely related to potato maturity. In Arabidopsis, this gene regulates flowering of *Arabidopsis thaliana* (Ma Da et al. 2014). Among the photoperiod, it regulated flowering pathways that currently have been discovered, a variety of components are E3 ubiquitin ligase target proteins and can be mediated by E3 ubiquitin ligase to achieve its ubiquitination degradation, thus affecting the photoperiod signals for flowering regulation and may affect photoreceptor stability, circadian clock function, and flowering regulator CO stability (Chen et al. 2011; Imaizumi et al. 2003; Takase et al. 2011; Ma Da, et al. 2014). Photoperiod has a great influence on potato plant growth, tuber formation, and development. Under long daylight, stems and leaves grow vigorously, a large number of stolons appear, but the tuber formation is delayed and the yield is decreased. Under short daylight, plant growth is normal, and the tuber production is faster. Assimilation products are transported to tubers faster, thus tuber yield is high. Early maturing varieties are sensitive to daylight, while late maturing varieties have to form tubers under short daylight conditions. Therefore, the signal transduction pathway of potato flowering and tuber formation may be similar to that of *Arabidopsis thaliana* flowing regulation. Thus, E3 ubiquitin ligase PUB14 may act as an intermediate mediator during potato flowering or tuber formation, regulating flowering and tuber formation. In addition, the auxin export vector gene may also be involved in potato maturity. Auxin participates in the regulation and control of many physiological and biochemical processes, such as root occurrence, photoreaction, apical dominance, flowers development, leaves and fruits shedding, and distribution of assimilation products. In the process of auxin transport, this protein, as a specific auxin output vector, can induce IAA passively flowing to the cell wall, then enter into next cell, thus forming polar transport. In the late growth and development of potato, this gene may be involved in assimilation product distribution as well as leaf and fruit shedding, thus plants show early maturation. However, whether these two genes are related to maturity needs further analysis and validation. In this study, we also examined the segregation of the maturity-related gene *StCDF1* (Kloosterman et al. 2013) in the population of this study, and the results showed that the late maturing parent Zhongshu 19 contained the gene *StCDF1.1* (late maturity related gene) and *StCDF1.2* (early maturity related gene), the early maturing parent Zhongshu 3 only contained *StCDF1.1,* both the parents do not contained *StCDF1.3* (early maturity related gene), and also there was no polymorphism in our segregation population. Moreover, we also test the *StCDF1.2* gene in 83 Chinese varieties, and the result showed that only 4 out of 36 early maturity varieties and 5 out of 47 late maturity varieties contained the early maturity related gene *StCDF1.2* (data not shown). Moreover, the alignment of *StCDF1* with the reference genome showed that the *StCDF1* was located in 4.537-4.542Mb, but the maturity locus in our study is located in 3.7-4.2Mb. Pedigree investigation showed that the *StCDF1.2* gene was cloned from CE3130 that should originated from phureja (Kloosterman et al., 2013), the early maturity genes of Zhongshu 3 maybe come from cv. Katahdin. All the above indicated that there should exist other different maturity-related genes in our maturity segregation population of this study from the gene *StCDF1.*

The tetraploid mapping software TetraploidMap (Hackett et al. 2007) used in this study only can identify three marker types. For a single dominant marker, the parental genotype is AOOO × OOOO, and the segregation ratio of progeny is 1:1. For double dominant markers, the parental genotypes are AAOO × OOOO and AOOO × AOOO, and the segregation ratios are 5:1 and 3:1. There are still two types of markers the software does not recognize, i.e. markers of parental genotypes AOOO × AAOO and AAOO × AAOO with the segregation ratios of 11:1 and 35:1. Therefore, among the 32 polymorphic markers selected from 152 SSR markers, four markers belonged to the 11:1 marker type, and the software did not recognize them. Therefore, only 28 polymorphic SSR markers could be used in the study to construct the map. Because the tetraploid mapping software limits the recognition of marker types, some markers are not available, and made the number of markers less than normal in the genetic map. Nevertheless, we have mapped the QTL for the early maturity trait in the physical region of 471 kb and analyzed the genes within this interval. Which provides a solid foundation for the cloning of the major gene or genes that control the early maturity trait.

## Acknowledgments

This work was supported by National Key R&D Program of China (2017YFD0101905), the National Natural Science Fund of China (31561143006, 31771860), China Agriculture Research System (CARS-9).

## Compliance with ethical standards

### Conflict of interest

Authors declare that they do not have conflict of interest.

## Table Captions

**Table 1** Information of the 50 primers used in the genetic map

**Table 2** Basic analysis of sequencing data

**Table 3** Makers associated with physiological maturity KWSig: the significant of Kruskal-Wallis test. AVSig: the significant of the analysis of variance. Mean (0):the mean when the maker is absent. Mean (1): the mean when the maker is present. SED: the stand error of difference between the means.

**Table 4** Gene annotation and location in the physical interval

**Table S1** Phenotypic data 2014-2017

Author contribution statement
GL and LJ conceived the project, provided idea and designed the whole experiments; XL and JZ conceived the project and performed the molecular marker development and analysis; JX contributed to the maturity phenotyping of the mapping population; SD performed the parents crossing of the mapping population; CB and JH performed the field management of potato materials. All authors read and approved the final manuscript.

